# Fast Whole-Genome Phylogeny of the COVID-19 Virus SARS-CoV-2 by Compression

**DOI:** 10.1101/2020.07.22.216242

**Authors:** Rudi L. Cilibrasi, Paul M.B. Vitányi

## Abstract

We analyze the whole genome phylogeny and taxonomy of the SARS-CoV-2 virus using compression. This is a new fast alignment-free method called the “normalized compression distance” (NCD) method. It discovers all effective similarities based on Kolmogorov complexity. The latter being incomputable we approximate it by a good compressor such as the modern zpaq. The results comprise that the SARS-CoV-2 virus is closest to the RaTG13 virus and similar to two bat SARS-like coronaviruses bat-SL-CoVZXC21 and bat-SL-CoVZC4. The similarity is quantified and compared with the same quantified similarities among the mtDNA of certain species. We treat the question whether Pangolins are involved in the SARS-CoV-2 virus. The compression method is simpler and possibly faster than any other whole genome method, which makes it the ideal tool to explore phylogeny.

## I. Introduction

In the 2019 and 2020 pandemic of the COVID-19 illness many studies use essentially two methods, alignment-based phylogenetic analyses e.g. [19], and an alignment-free machine learning approach [23]. These pointed to the origin of the SARS-CoV-2 virus which causes the COVID-19 pandemic as being from bats. It is thought to belong to lineage B (Sarbecovirus) of Betacoronavirus. From phylogenetic analysis and genome organization it was identified as a SARS-like coronavirus, and to have the highest similarity to the SARS bat coronavirus RaTG13 [19] and similar to two bat SARS-like coronaviruses bat-SL-CoVZXC21 and bat-SL-CoVZC45.

Alignment methods are generally used but have many drawbacks which make alignment-free methods more attractive [28], [27], [30]. The purpose of this study is to introduce a new alignment-free method based on lossless compression of the whole genome sequences of base pairs of the involved viruses, the *normalized compression distance* (NCD) method. We use it here to identify the 60 closest viruses to the SARS-CoV-2 virus and to determine their relations (taxonomy) in a phylogeny tree. With this method we can actually quantify these relations and compare them to similar relations between the mtDNA’s of mammal species in order to gain an intuition as to what they mean.

We computed the NCD’s of 6,751+15,409=22,160 virus pairs plus the NCD’s used for Figures 1 and 2. Altogether, this took about 5–10 hours in a combined run on a home desktop computer (a mini-computer called Meerkat from a Linux computer company called System76). It uses less than 2 seconds per pair which comes to more than 2000 pairs per hour. The same program can be reused for different phylogenies and questions. This is possibly the easiest and fastest method to establish whole genome phylogeny.

**Fig. 1.**
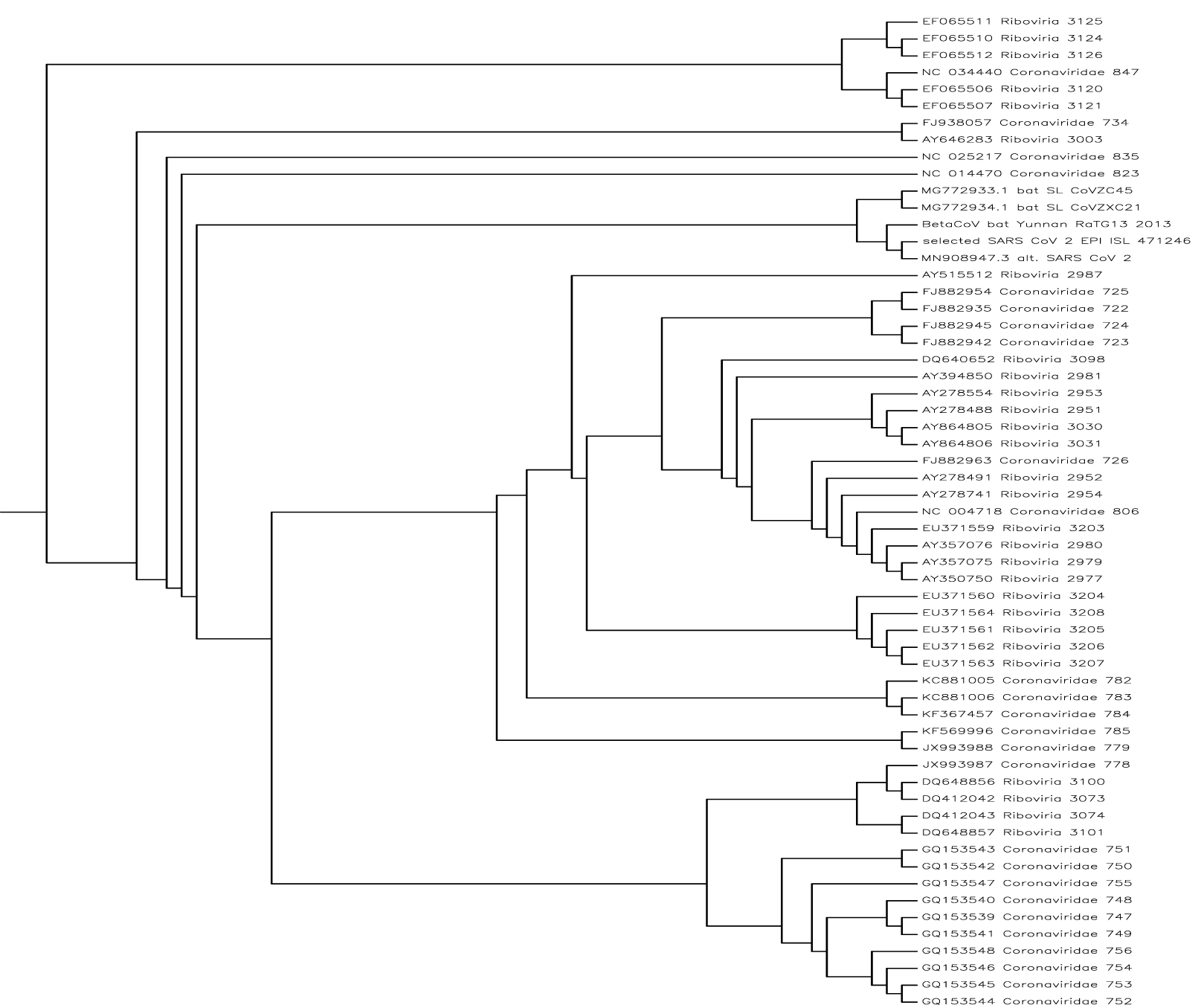
The evolutionary directed binary tree built from the 60 virus sequences in Table I. These viruses have the least NCD distance with the SARS-CoV-2 virus. *S*(*T*) = 0.998948. We use human mtDNA as outgroup; it is about the same size as the viruses and it can be assumed to be completely different. Where it joins the tree there is the root. The labeling of the items is as follows. All sequences are labeled as they occur in the data of [23] together with their registration code. The most interesting are the 11th to 15th (inclusive) sequences of the tree from the top of the page. The 13th to 15th virus sequences (inclusive) are the most interesting for us. The 13th is EPI_ISL_402131 which is the bat/Yunnan/RaTG13/2013 that is the RaTG13 Bat Coronavirus sampled in Yunnan in 2013. The 14th label is selected SARS-CoV-2 virus which occurs 105 times in the virus database of GISAID. It is the top one in the sorted list Table I with NCD of about 0 with respect to the selected SARS-CoV-2 virus, that is, it is the same virus EPI_ISL_428253. The 15th label is MN908947 which is a a SARS-CoV-2 virus Wuhan Hu-1 from the Wuhan Seafood market collected Dec 2019, submitted 05-JAN-2020, and reported in *Nature*, 579(7798), 265–269, 270–273 (2020). This is the only SARS-CoV-2 virus in the 6,751 sequences obtained from the authors of [23]. The 11th and 12th labels are the CoVZC45 Bat Coronavirus and the CoVZXC21 Bat Coronavirus. Numbers 11, 12, 13, 14, and 15 have the least NCD distances to the selected SARS-CoV-2 virus. Here are the NCD’s of the selected SARS-CoV-2 virus against 59 others: selected_SARS_CoV_2_EPI_ISL_471246 0.932533 0.994986 0.918724 0.919072 0.925802 0.919182 0.918691 0.995086 0.91954 0.918829 0.444846 0.926681 0.926013 0.995086 0.952228 0.919486 0.91801 0.995078 0.918531 0.918381 0.919117 0.918257 0.92615 0.788416 0.925982 0.994546 0.00362117 0.917563 0.923045 0.92615 0.918658 0.918597 0.920486 0.919221 0.917493 0.91993 0.92541 0.925844 0.921053 0.994986 0.918945 0.925951 0.923151 0.995078 0.0111034 0.931577 0.918497 0.918553 0.995078 0.925706 0.791082 0.919244 0.92613 0.997632 0.994897 0.918447 0.918796 0.918605 0.918669 0.918565. The entire 60 × 60 NCD distance matrix underlying this tree is too large to display but available from the authors on request.

**TABLE I.**
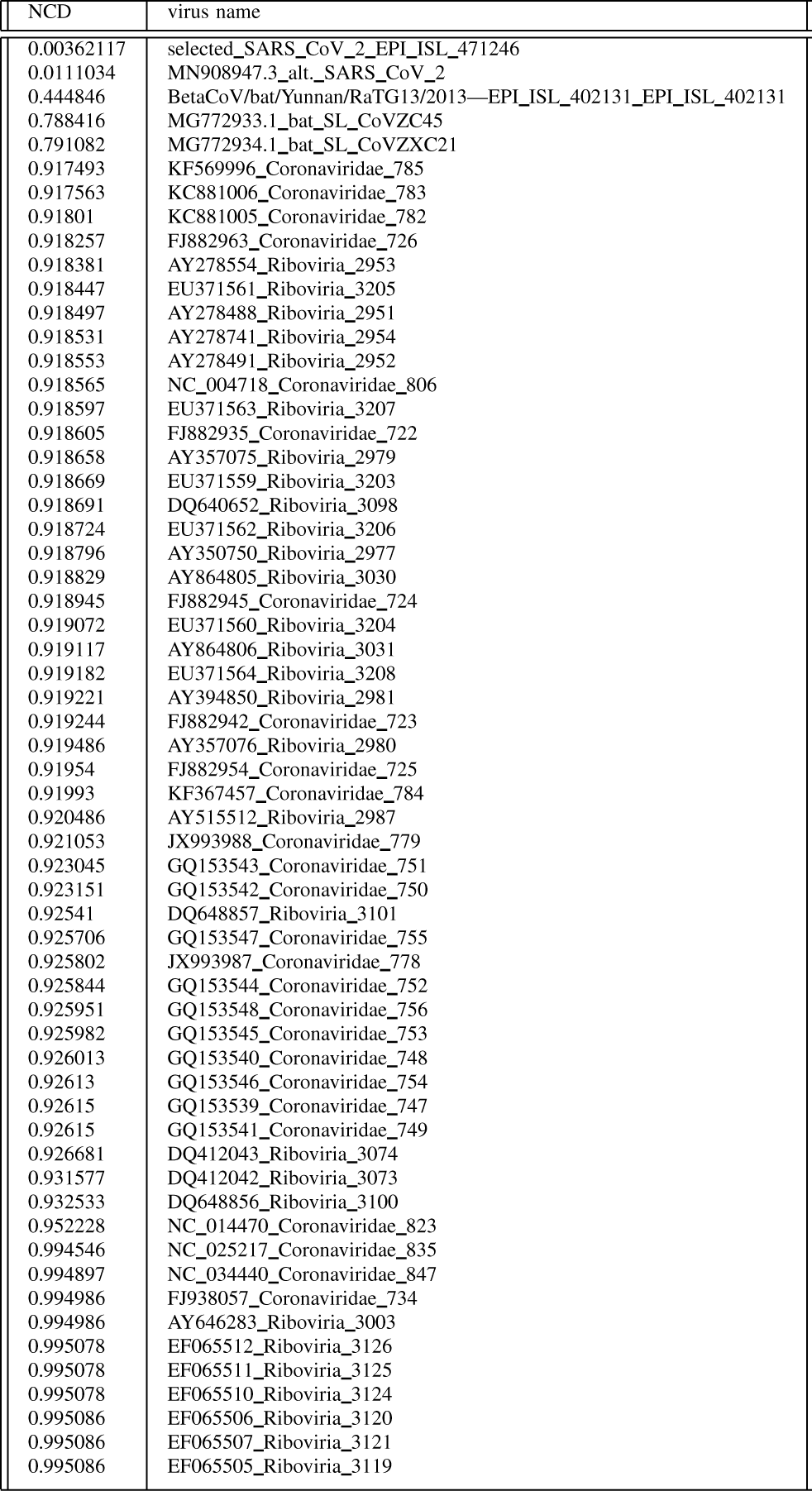
First 60 items in the sorted list of used 6,751 virus sequences.

**TABLE II.**
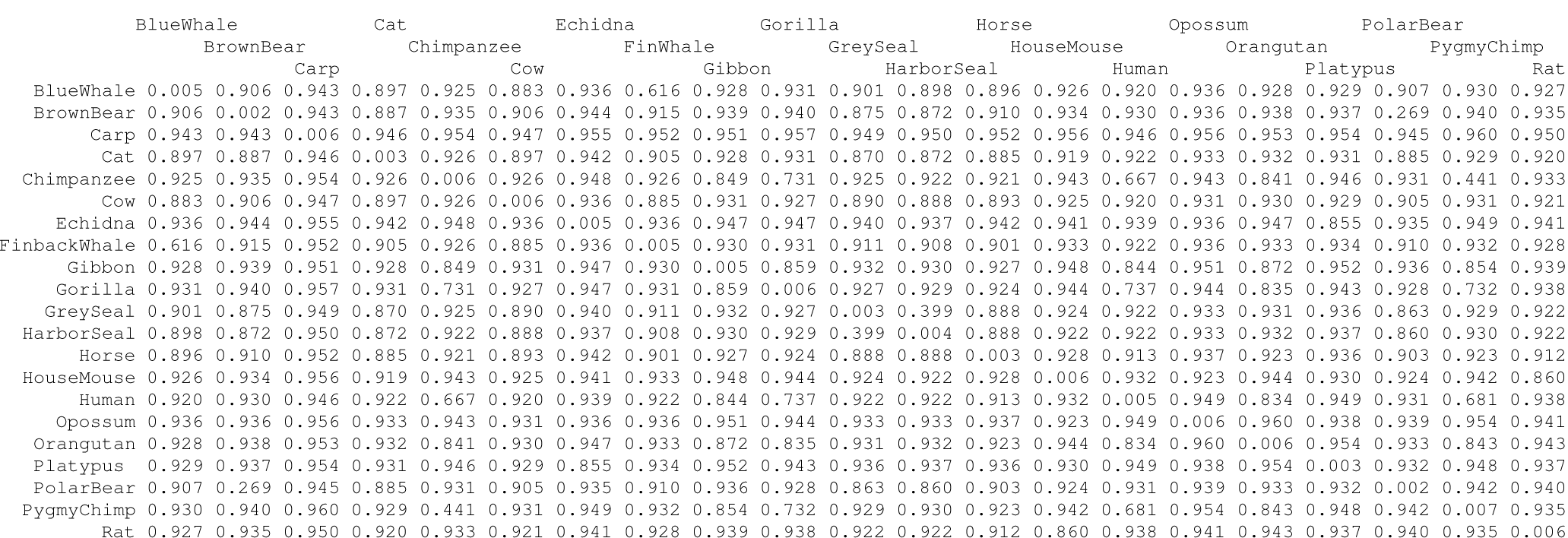
Distance matrix of pairwise NCD. For display purpose, we have truncated the original entries from 15 decimals to 3 decimals precision.

**Fig. 2.**
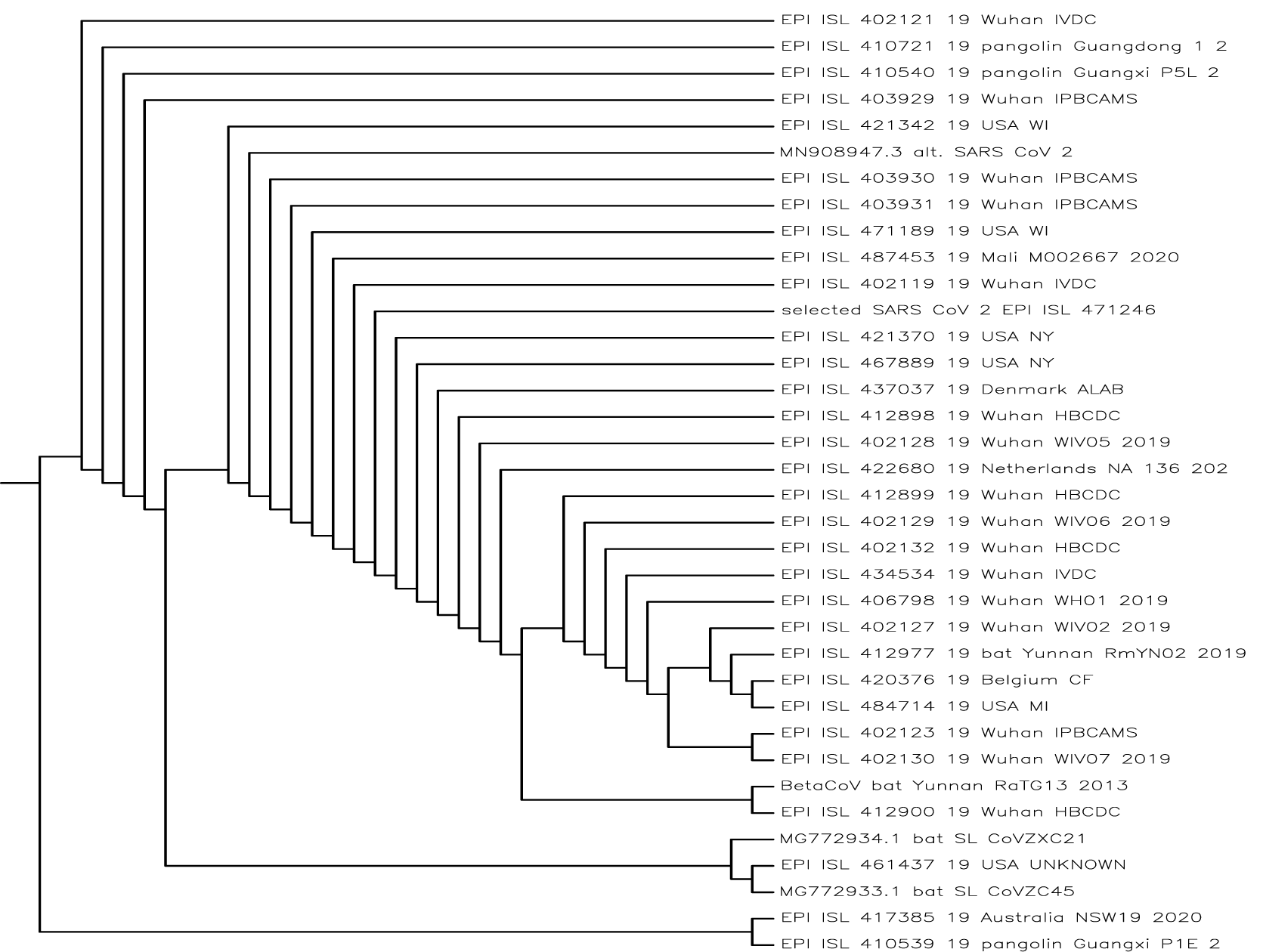
The evolutionary directed binary tree built from 37 virus sequences with the human mtDNA to determine the root. It contains the selected SARS-CoV-2 sequence and all the GISAID sequences ending in /2017, /2018, and /2019 (this includes the three pangolins). Added are a dozen close sequences from all GISAID sequences and a dozen close sequences from the machine learning approach study [23] data. The ladders in the directed binary tree usually accomodate more than two outgoing branches. Here are the NCD’s of the selected SARS-CoV-2 virus against the other viruses: selected_SARS_CoV_2_EPI_ISL_471246 0.010421 0.00417827 0.444846 0.0412256 0.0130447 0.0111034 0.435097 0.00375992 0.788416 0.00362117 0.0106989 0.738314 0.00986522 0.0122137 2.04891e-08 0.873888 0.873228 0.0115198 0.0117974 0.004039 0.00389972 0.0111034 0.0116699 0.00417711 0.0116699 0.0111034 0.0116699 0.0123525 0.791082 0.997632 0.00417827 0.00417827 0.0105702 0.00417827 0.00417827 0.004039 0.0101417. The entire 37 × 37 NCD distance matrix is too large to display but available from the authors on request.

## II. Materials and Method

We first obtain viruses (in this case at least 6,500 of them) we want to compare the SARS-CoV-2 virus to, without duplicates, partially sequenced viruses, and SARS-CoV-2 viruses. RNA sequences have A,C,G,U and DNA have A,C,G,T base pairs, but since SARS-CoV-2 is an RNA virus but all sequences considered here have base pairs A,C,G,T, technically the sequences are cDNA (complementary DNA) of the RNA sequence in the virus itself. We select a SARS-CoV-2 virus from the many, at least 15,500, examples available. Compute for each virus we want to compare with the selected SARS-CoV-2 virus its NCD distance with the selected SARS-CoV-2 virus. For an explanation of the NCD see the Appendix. Subsequently we order the resulting NCD’s from the smallest to the largest. The virus causing the smallest NCD distance with the SARS-CoV-2 virus is the most similar to that virus. We next take the 60 viruses which have the least NCD’s with the selected SARS-CoV-2 virus and compute the phylogeny of those viruses and the SARS-CoV-2 virus. Next we compare 37 viruses to determine the relation of the SARS-CoV-2 virus with Pangolins.

## III. Data and Data Cleaning

We downloaded the data broadly in two parts. The original input data are stored by Rudi Cilibrasi at:

-rw-rw-r– 1 rudi rudi 2.0G Jul 17 15:46 incoming/gisaid_hcov-19_2020_07_17_22.fasta

-rw-rw-r– 1 rudi rudi 29M Jul 17 15:45 incoming/gurjit-data.zip

We downloaded from the GISAID [11] database on July 17th, 2020 in total 66,899 sequences. A part of at least 99.9% of the GISAID data are SARS-CoV-2 viruses. The remainder are not technically SARS-CoV-2 viruses but are related and of interest. One of the authors of the machine learning approach study [23] was so kind as to supply us with the 6,751 sequences used in that paper. There is one SARS-CoV-2 virus among them. Thus, at least 99.9% of the data is not SARS-CoV-2 viruses but other viruses. Therefore we can compare our results about the phylogeny of SARS-CoV-2 directly with those of [23].

We removed a single sequence with a special exception part in the code because it had a bad name: gisaid_hcov-19_2020_07_17_22.fasta.fai hCoV-19 29868 1713295574 80 82. That name is too short to be useful: “hCoV-19.” Altogether therefore we imported 73,649 sequences total into the raw sequence database. Each viral sequence is an RNA sequence and seems to be around 30,000 RNA base pairs (A,T,G,C) in size. The total size of all sequence data together is in the order of two Gigabyte. Of the sequences initially downloaded from GISAID, we applied a lowercase transformation to each to reduce pointless variability. After that, we computed a histogram of all the characters in the sequence and counted the size of each group. Many sequences contained the base pairs A, C, G, N, T or other letters. The letters A,C,G, and T signify the basic DNA nucleotides. the meaning of ‘N’ and the other letters involved can be found on a FASTA file format reference webpage. For example, “N” means “unknown nucleotide residue.” We retained the deduplicated unique 15,578 sequences with the known nucleotides A,C,G, and T from the GISAID download. The 6,751 sequences obtained from the authors of [23] were over the letters A,C,G, and T already.

As a representative of the SARS-CoV-2 virus we selected the most common one in the dataset as the basis for the NCD against all others. It appeared in the sorted virus list with 105 multiples. In Section IV it appears at the top with an NCD of 0.003621 and has the official name of gisaid_hcov-19_2020_07_17_22.fasta<hCoV-19/USA/WI-WSLH-200082/2020—EPI_ISL_471246—2020-04-08.

We then looked at whether there was much variation among the SARS-CoV-2 viruses themselves since this may invalidate the NCD distance between the inspected viruses and the selected SARS-CoV-2 virus. We retained the viruses in the list after deduplication and filtering for A,C,G,T. The worst NCD against the selected SARS-CoV-2 virus was 0.874027 namely gisaid_hcov-19 2020 07 17 22.fasta <hCoV-19/pangolin/Guangxi/P1E/2017EPI ISL 4105392017 from a Pangolin. Removing that one sequence from the list we got a worst NCD of 0.873367 also from a Pangolin in 2017.

(To clarify why there is the discrepancy below in that we obtain 15,578 imported sequences but when counting lose 100–200 in the next phase: There are some exact name duplicates in the GISAID data. When the “imported sequence” count is reported, then we count identically named identical sequences separately because they did both get imported (but one would overwrite the other). When counting the different names for sequences without /2017, /2018, /2019 in the name, these exact-name duplicates would collapse into a single one causing just one count.)

Initially there are 15,430 sequences from GISAID that contain “hCov” in the name. We then removed all sequences that contained /2017 in the name. After this we were left with 15,428 sequences and obtained a worst NCD of 0.738175 also from a Pangolin.

Removing all sequences that contained /2017, /2018, or /2019 in the name, left 15,409 viruses in the list. The worst-case NCD across all the remaining sequences of the SARS-CoV-2 virus to the selected SARS-CoV-2 virus is 0.044986 and the average is 0.009879. This shows that the 15,428 SARS-CoV-2 viruses retrieved from GISAID on July 17 in 2020 all have sequences that hardly differ from one another. The worst-case sequence [20] can be found at gisaid_hcov-19_2020_07_17_22.fasta<hCoV-19/USA/OH-OSUP0019/2020—EPI_ISL_427291—2020-03-31 and its registration code is EPI_ISL_427291 [20].

With respect to the figures: Figures 1 and 2 were done with the PHYLIP (PHYLogeny Inference Package) [29] and Figure 3 with the Complearn Package [5]. The programming is easy. Essentially one only has to compute Equation A.4 for all pairs of viruses to be compared. The comparisons between the NCD’s of viruses and the NCD’s of the mtDNA’s of species are made on basis of Table II. Since the NCD is a metric (satisfies the triangle property) [6, Theorem 6.2] the above variation of NCD’s of different SARS-CoV-2 viruses will not unduly upset the greater NCD distances (only by ±0.044986) between the other viruses (the 6,751 non-SARS-CoV-2 viruses) and the selected SARS-CoV-2 virus. The analytic software which was used is available at Github source code URI: https://github.com/rudi-cilibrasi/ncd-covid

**Fig. 3.**
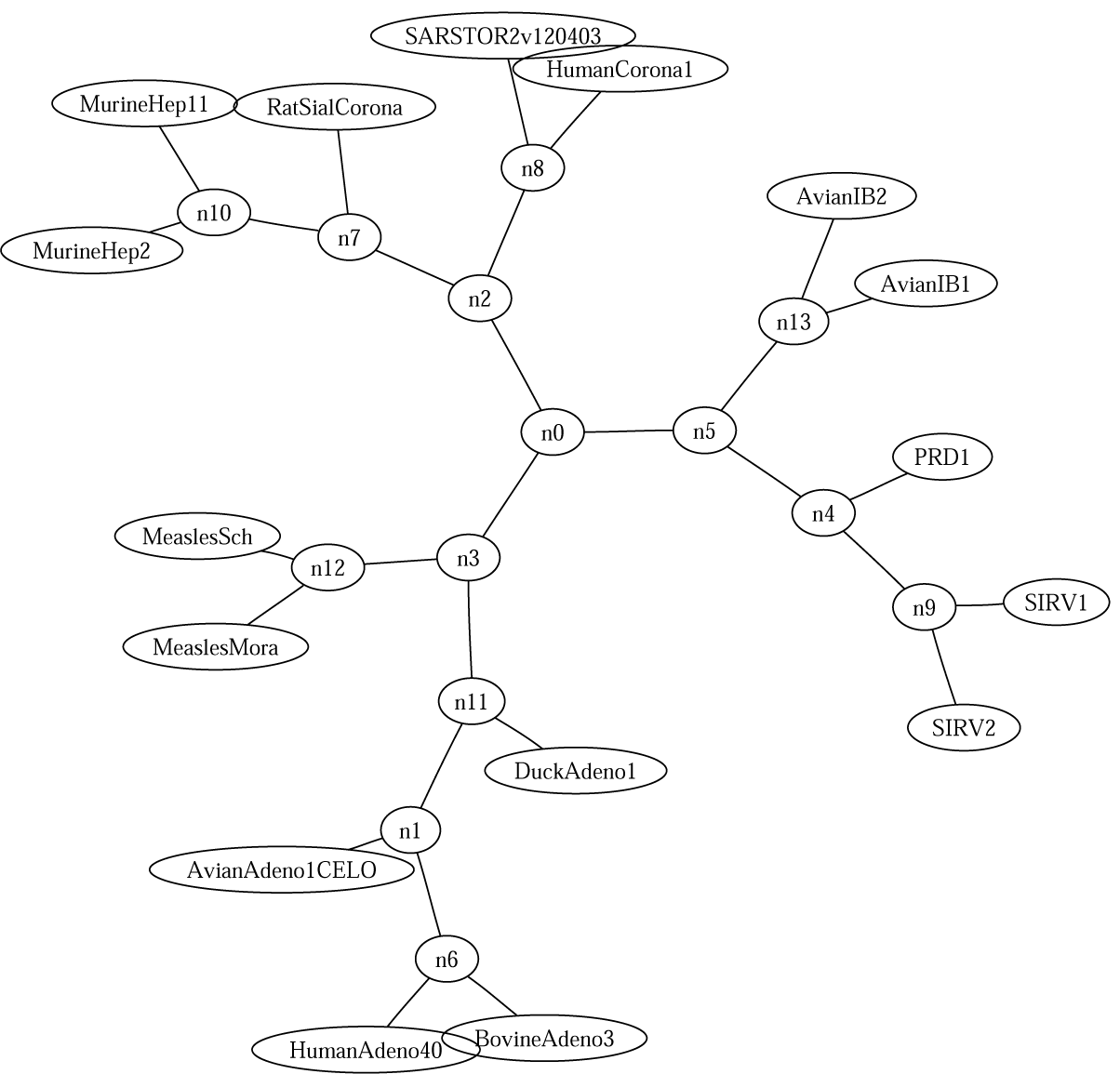
SARS virus among other viruses, *S*(*T*) = 0.988. Explanation text in nodes: AvianAdeno1CELO.inp: Fowl adenovirus 1; AvianIB1.inp: Avian infectious bronchitis virus (strain Beaudette US); AvianIB2.inp: Avian infectious bronchitis virus (strain Beaudette CK); BovineAdeno3.inp: Bovine adenovirus 3; DuckAdeno1.inp: Duck adenovirus 1; HumanAdeno40.inp: Human adenovirus type 40; HumanCorona1.inp: Human coronavirus 229E; MeaslesMora.inp: Measles virus strain Moraten; MeaslesSch.inp: Measles virus strain Schwarz; MurineHep11.inp: Murine hepatitis virus strain ML-11; MurineHep2.inp: Murine hepatitis virus strain 2; PRD1.inp: Enterobacteria phage PRD1; RatSialCorona.inp: Rat sialodacryoadenitis coronavirus; SARS.inp: SARS TOR2v120403; SIRV1.inp: Sulfolobus virus SIRV-1; SIRV2.inp: Sulfolobus virus SIRV-2.

## IV. Results

In the sorted list of NCD’s between the selected SARS-CoV-2 virus and the 6,751 other viruses the selected SARS-CoV-2 virus appears on the top of the list with NCD toward itself of 0.00362117 (should be 0 but the computation has a small error margin). The next virus is the only SARS-CoV-2 virus in the database of 6,751 virus sequences (not the selected one) with NCD=0.0111034. This gives confirmation that the NCD’s in the list are accurately calculated since the number is so very low. The code is MN908947.3. It is isolate Wuhan-Hu-1, complete genome GenBank: MN908947.3 of 29903 bp ss-RNA linear VRL 18-MAR-2020 of the family Viruses; Riboviria; Orthornavirae; Pisuviricota; Pisoniviricetes; Nidovirales; Cornidovirineae; Coronaviridae; Orthocoronavirinae; Betacoronavirus; Sarbecovirus.

The following virus has an NCD of 0.444846 with the selected SARS-CoV-2 virus and is the closest (apart from the above SARS-CoV-2 virus). It is Sarbecovirus/EPI_ISL_402131.fasta/ <BetaCoV/bat/Yunnan/RaTG13/2013—EPI_ISL_402131. The first part is the classification of the subfamily of viruses, the code EPI_ISL_402131 is that of the virus itself which one can use with Google to obtain further information.

In this case the virus is sampled from a bat in Yunnan Province in in the PRC (China) in 2013 and is 29855 bp RNA linear VRL 24-MAR-2020, and its final registration code is MN996532. It is known as Bat coronavirus RaTG13 of the family Viruses; Riboviria; Orthornavirae; Pisuviricota; Pisoniviricetes; Nidovirales; Cornidovirineae; Coronaviridae; Orthocoronavirinae; Betacoronavirus; Sarbecovirus. The virus is found in the Rhinolophus affinis, a medium-size Asian bat of the Yunnan Province (China). The human coronavirus genome shares at least 96.2% of its identity with its bat relative, while its similarity rate with the human strain of the SARS virus (Severe Acute Respiratory Syndrome) is much lower, only 80.3% [8]. The NCD distance between the selected RSA-CoV-2 virus and this virus is about the same as that between the mtDNA’s of the Chimpansee and the PigmyChimpansee according to the Table II.

The next three viruses have respectively NCD=0.788416 for Coronaviridae/CoVZC45.fasta/ < MG772933.1, and NCD=0.791082 for Coronaviridae/CoVZXC21.fasta/ < MG772934.1 and at a larger distance NCD=0.917493 for Coronaviridae/Coronaviridae_783.fasta/ < KC881006 Here the part “fasta/ < KC881006” means that the registration code is KC881086.

The first of these two viruses is the Bat SARS-like coronavirus isolate bat-SL-CoVZC45, complete genome at 29802 bp RNA linear VRL 05-FEB-2020 of the above family of viruses. Its NCD with the selected SARS-CoV-2 virus is slightly larger than the mtDNA distance between the Human and the Gorilla at 0.737 and slightly lower than the mtDNA distance between the Human and the Orangutan at 0.834.

The second of these two viruses is the Bat SARS-like coronavirus isolate bat-SL-CoVZXC21, complete genome at 29732 bp RNA linear VRL 05-FEB-2020 of the same family. The same comparison of the NCD distance between this virus and the selected SARS-CoV-2 virus with the NCD of mtDNA’s between (the same) species holds also for this second virus.

The third virus in the list is the Bat SARS-like coronavirus Rs3367, complete genome at 29792 bp again of the same family. Comparing the NCD between this virus and the selected SARS-CoV-2 virus yields that it is slightly smaller than the NCD between the Human mtDNA and the Blue Whale mtDNA at 0.920 and slightly larger than between the mtDNA of the Finback Whale and the mtDNA of the Brown Bear at 0.915.

We can conclude that the SARS-CoV-2 virus is likely from the family Viruses; Riboviria; Orthornavirae; Pisuviricota; Pisoniviricetes; Nidovirales; Cornidovirineae; Coronaviridae; Orthocoronavirinae; Betacoronavirus; Sarbecovirus.

### A. The Pangolin connection

As we saw in determining the worst-case NCD between the selected SARS-CoV-2 virus and other GISAID SARS-CoV-2 database viruses the database is littered with Pangolin viruses from 2017, 2018, and 2019. Several studies e.g. [16], [31] hold that while the SARS-CoV-2 virus probably originates from bats it may have been transmitted to another animal and/or recombined with a virus there and transmitted zoonotic to humans. The other animal is most often identified as the Pangolin. The compression method shows that the NCD’s between the Pangolin SARS-CoV-2 virus and the human SARS-CoV-2 virus are far apart. However, they are not farther apart than 0.738175 to 0.874027. This is between the NCD distance of the mtDNA of a Human to a Gorilla up to the mtDNA of a Greyseal to a Bluewhale. The bat-SL-CoVZXC21 and bat-SL-CoVZC45 viruses are at NCD=0.791082 and NCD=0.788416. That is in between the mtDNA NCD distance of a Human versus a Gorilla at 0.737 and the mtDNA distance of a Human versus an Orangutan at 0.834. Thus the Pangolin and bat origins are possible for the SARS-CoV-2 virus while the bat origin is more likely and modification by the Pangolin is a possibility from these data. But the RaTG13 virus has an NCD=0.444846 distance with the human SARS-CoV-2 virus that is close to one half of the Pangolin distance. As noted before, this is comparable with the NCD distance between the Chimpansee and the PigmyChimpansee according to the Table II. Also in the tree of Figure 2 the Pangolin viruses are generally far from the selected SARS-CoV-2 virus. Hence the hypothesis that the Pangolin species is an intermediary between the bat viruses above and the human SARS-CoV-2 virus is perhaps unlikely.

## V. Discussion

Previously most studies into the COVID-19 pandemic suggested that the virus involved originated from bats. Bats are a known reservoir of viruses that can zoonotic transmit to humans [18]. Virtually all those mentioned studies use alignment-based methods. Analyzing the involved SARS-CoV-2 virus with the NCD is in this setting a novel alignment-free method based on compression. It places the RaTG13 bat virus the closest to the SARS-CoV-2 virus followed by the two SARS-like coronaviruses bat-SL-CoVZXC21 and bat-SL-CoVZC4. The similarity is quantified and compared with the same quantified similarities among the mtDNA of certain species. A possible involved other animal is the Pangolin. The method used here, the normalized compression distance (NCD) method is based on Kolmogorov complexity analysis and compression [15], [6]. It is domain-independent and requires no parameters to be set, apart from the used compression algorithm. Earlier studies using alignment-based methods have suggested that the the SARS-CoV-2 virus originated from bats before being zoonotical transferred to humans. The machine-learning approach, an alignment-free method, in [23] came to the same conclusion. The current, completely different, alignment-free method with the results we present here, confirms this conclusion. Hence there is little doubt by now that the virus involved originates from bats with as runner-up the Pangolin.

## VI. Conclusion

We provide a new alignment-free method based on compression to determine the phylogeny and taxonomy of the SARS-CoV-2 virus (the virus causing the COVID-19 pandemic). The method is based on compression, remarkably simple, very fast, and quantifies a distance to a number between 0 (identical) and 1 (totally different). It uses only the unprocessed viral DNA/RNA sequences. In the Appendix it is briefly explained and illustrated by examples of mammal species via mtDNA sequences and the phylogeny and taxonomy of the SARS virus. To compare it with another alignment-free method based on machine learning and other techniques [23] we used the same database of over 6000 unique viral sequences. To select the sequence of the SARS-CoV-2 virus against which the unique viral sequences are compared we use a database of over 15,000 unique SARS-CoV-2 viruses. Obtaining similar results as the earlier studies, and confirming the generally believed hypothesis, this method is less complicated than the previous methods. It yields quantitative evidence that can be compared with similar distances among the mtDNA of familiar mammals. Since the method is uncomplicated and very fast it is useful as an exploratory investigation into the phylogeny and taxonomy of viruses of new epidemic outbreaks.

## Appendix

In 1936 A.M. Turing [25] defined the hypothetical “Turing machine” whose computations are intended to give a formal definition of the intuitive notion of computability in the discrete domain. These Turing machines compute integer functions, the *computable* functions. By using pairs of integers for the arguments and values we can extend computable functions to functions with rational arguments and/or values.

### A. Kolmogorov Complexity

Informally, the Kolmogorov complexity of a string is the length of a shortest string from which the original string can be losslessly reconstructed by an effective general-purpose computer such as a particular universal Turing machine *U*. Hence it constitutes a lower bound on how far a lossless compression program can compress. For details see the text [17]. In this paper we require that the set of programs of *U* is prefix free (no program is a proper prefix of another program), that is, we deal with the *prefix Kolmogorov complexity*. We call *U* the *reference universal prefix machine*. Formally, the *conditional prefix Kolmogorov complexity K*(*x*|*y*) is the length of a shortest input *z* such that the reference universal prefix machine *U* on input *z* with auxiliary information *y* outputs *x*. The *unconditional prefix Kolmogorov complexity K*(*x*) is defined by *K*(*x*|*ϵ*) where *ϵ* is the empty word of length 0.. The functions *K*(·) and *K*(·|·), though defined in terms of a particular machine model, are machine-independent up to an additive constant and acquire an asymptotically universal and absolute character through Church’s thesis, and from the ability of universal machines to simulate one another and execute any effective process.

The Kolmogorov complexity of an *individual finite object* was introduced by Kolmogorov [12] as an absolute and objective quantification of the amount of information in it. It is sometimes confused with the information theory of Shannon [21], which deals with *average* information *to communicate* objects produced by a *random source*. They are quite different.

### B. Information Distance

The *information distance D*(*x, y*) between strings *x* and *y* is defined as

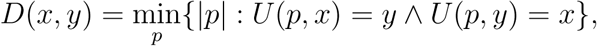

where *U* is the reference universal prefix machine above. Like the Kolmogorov complexity *K*, the distance function *D* is upper semicomputable. Define

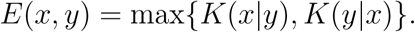

In [1] it is shown that the function *E* is upper semicomputable, (Here and elsewhere in this paper “logarithm” or “log” refers to the binary logarithm.) *D*(*x, y*) = *E*(*x, y*)+*O*(log *E*(*x, y*)), the function *E* is a metric (more precisely, that it satisfies the metric (in)equalities up to a constant), and that *E* is minimal (up to a constant) among all upper semicomputable distance functions *D*′ satisfying the normalization conditions ∑_*y*:*y*≠*x*_ 2^−*D*′(*x,y*)^ and ∑_*x*:*x*≠*y*_ 2^−*D*′(*x,y*)^ (to exclude bogus distances which state, for example, that every *y* is in distance 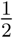 of a given *x*). We call this metric *E universal*.

Thus, for every pair of finite files *x, y* we have that *E*(*x, y*) is at least as small as the smallest *D*′(*x, y*). This means that *E*(*x, y*) is at least as small as the distance engendered by the dominant feature shared between *x* and *y*. The *normalized information distance* (NID) is defined by

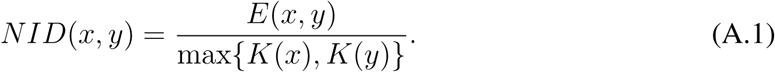

It is straightforward that 0 ≤ *NID*(*x, y*) ≤ 1 (up to an *O*(1*/* max{*K*(*x*), *K*(*y*)}) additive term). It is is a metric [15] (and so is the NCD we will meet in (A.4) see [6]. As an aside, a nonoptimal precursor to the NID/NCD was given in [14].) Since by the symmetry of information law [10] or see [17],

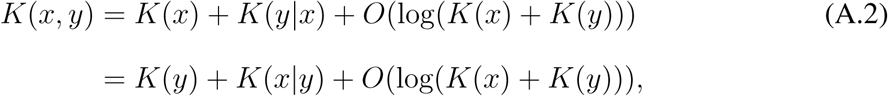

rewriting the NID using (A.2) yields

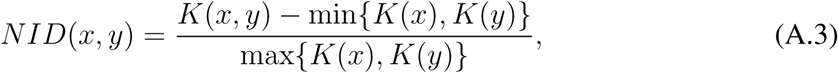

up to some small aditive terms that we ignore. For more details on this derivation see [17] or [26].

In this way, in [1], [15] we and others developed theoretical approaches to the similarity of finite objects. We proved that these theories based on Kolmogorov complexity are perfect. By approximating the Kolmogorov complexities involved by real-world compressors we transformed these theoretical notions into applications that work better than we could expect [6], [7]. It turns out that on natural data the above process gives adequate results. The resulting similarity measure is a parameter-free and alignment-free method. (In bioinformatics the computation of the similarity between genetic strings commonly involves the so-called “alignment method.” This method incurs often a high or even forbidding computational cost. for certain problems biologists look for alignment-free methods.) It is a *non feature-based similarity*. That is, it captures *every effective distance*: effective versions of Hamming distance, Euclidean distance, edit distances, alignment distance, Lempel-Ziv distance, and so on. It can simultaneously detect *all* similarities between pieces that other effective distances can detect separately.

Let us give an intuitive explanation. Two objects are deemed close if we can significantly “compress” one given the information in the other, the intuition being that if two pieces are more similar, then we can more succinctly describe one given the other. The NID discovers all effective similarities in the sense that if two objects are close according to *some* effective similarity then they are also close according to the NID. One of the advantages of this similarity is that it works for noisy objects [4]. It is a quantity between 0 (identical) and 1 (completely dissimilar).

### C. Normalized Compression Distance

Unfortunately, the universality of the NID comes at the price of incomputability. In fact, it is not even semicomputable and there is no semicomputable function at a computable distance of it [24]. One uses real data-compression programs to approximate the Kolmogorov complexity. The length of the compressed version of a finite object is obviously computable. Usually the computation process is fast. For the natural data we are dealing with we assume that the length of the compressed version is not too far from its Kolmogorov complexity. We substitute the Kolmogorov complexity in the NID by its approximation. If *Z* is a compressor and we use *Z*(*x*) to denote the length of the compressed version of a string *x*, then we arrive at the *Normalized Compression Distance*:

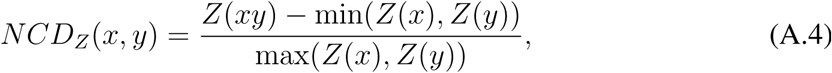

where we have replaced the pair (*x, y*) in the formula by the concatenation *xy* (file *y* appended to file *x*) and we ignore logarithmic terms in the numerator and denominator, see [6].

#### Remark 1

In [6] there are axioms to capture the real-world setting, and show that (A.4) approximates optimality. Actually, the NCD is a family of compression functions parameterized by the given data compressor *Z*. Common compressors are gzip (a Lempel-Ziv compressor with small window size of 32kB), bzip2 (a block-sorting compressor based on the Burrows-Wheeler transform with a larger window size of 256kB), and PPMZ (prediction by partial matching (PPM) which is an adaptive statistical data compression technique based on context modeling and prediction [9].

The objects being compared for similarity must fit in, say, one-half the window size. Gzip is by far the poorest compressor while PPMZ the best (although slowest) in this lineup. For example, the ideal compressor *Z* takes care that *NCD*_*Z*_(*x, x*) equals 0. With *Z* =gzip usually it is between 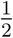 and 1 (very bad). With *Z* =bzip2 it is lower but nowhere near 0, and *NCD*_*PPMZ*_(*x, x*) in the genomic experiment of Table II below was between 0.002 and 0.006. For more experimental evidence see [3]. ●

#### Remark 2

Because of the normalization it does not matter for the NCD whether the length of data set *x* is different from the length of *y*. In practice, this difference should not be too great. ●

To visualise the *n*^2^ pairwise distances between *n* data we construct a dendrogram. To do so we take the *n* × *n* distance matrix as input, and construct a dendrogram with the *n* objects as leaves (so the dendrogram contains *n* external nodes or leaves and *n* − 2 internal nodes like in the figures below). We assume *n* ≥ 4. The resulting dendrogram models the distance matrix as good as possible qualitatively. If the distance between object *o*_1_ and object *o*_2_ is smaller that between *o*_1_ and *o*_3_, then the shortest path in the dendrogram between *o*_1_ and *o*_2_ has at most as many edges as the shortest path between *o*_1_ and *o*_3_ (equal if *o*_2_ and *o*_3_ are sibling nodes.) Thus, the edges themselves have no length and the dendrogram represents the partial order induced by the distance matrix. The *S*(*T*) value in Figures 1 and 3 (with *S*(*T*) = 1 is as good as possible) tells how well the tree represents the distance matrix [6]. For details see the cited reference. The method is available as an open-source software tool [5].

### D. Phylogeny

A DNA sequence is a finite string over a 4-letter alphabet {*A, C, G, T*}. We used the mitochondrial genomes (mtDNA) of 21 mammals, each of at most 18,000 base pairs, obtained from the GenBank Database. Hence, the mtDNA of every species involved is a string of at most 36,000 bits. Since we use the entire mtDNA of every species involved we do “whole-genome” phylogeny.

Whole genome phylogeny is usually only feasible with alignment-free methods, like the NCD method. This type of phylogeny is often computationally forbidding for the usual alignment methods used in bioinformatics. Moreover, gene areas move easily over the genome to other places again making the use of these methods impossible or hard. Hence it is more usual in bioinformatics to select a particular gene from the genome of each species. This particular gene should not evolve too fast, like the gene coding for insulin. Mutations here are usually fatal for the individual concerned. Thus, biologists feel that comparing these genes of species gives trustworthy information about the evolution of species. This may be called “gene tree phylogeny.” See [22]. However, using different genes may result in different trees [2]. But using the whole genome gives a single tree. For the 21 original species used we do not give the Latin names; they can be found in [6].

For every pair of mitochondrial genome sequences *x* and *y*, we evaluated the formula in Equation A.4 using a good compressor like PPMZ The resulting distances are the entries in an 21 × 21 distance matrix Table II. Constructing a phylogeny tree from the distance matrix, for example using our quartet tree method [7] or PHILIP [29] as tree-reconstruction software, gives the corresponding tree.

### E. SARS Virus

We clustered the SARS virus directly after its sequenced genome was made publicly available, in relation to potential similar viruses. The 15 virus genomes were downloaded from The Universal Virus Database of the International Committee on Taxonomy of Viruses, available on Internet. The SARS virus was downloaded from Canada’s Michael Smith Genome Sciences Centre which had the first public SARS Coronovirus draft whole genome assembly available for download (SARS TOR2 draft genome assembly 120403). The NCD distance matrix was computed using the compressor bzip2. The entire computation took only a couple of minutes. The relations in Figure 3 are very similar to the definitive tree based on medical-macrobio-genomics analysis, appearing later in the New England Journal of Medicine [13].

We depicted the figure in the ternary tree style, rather than the genomics-dendrogram style, since the former is more precise for visual inspection of proximity relations.

## Acknowledgement

The authors thank Gurjit Randhawa for the database of viruses used in [23].

## Author Contributions

**Conceptualization:** Paul Vitányi.

**Data curation:** Rudi Cilibrasi.

**Formal analysis:** Paul Vitányi.

**Methodology:** Rudi Cilibrasi, Paul Vitányi.

**Resources:** Rudi Cilibrasi.

**Software:** Rudi Cilibrasi.

**Supervision:** Paul Vitányi.

**Writing:** Paul Vitányi.

